# Corals sustain growth but not skeletal density across the Florida Keys Reef Tract despite ongoing warming

**DOI:** 10.1101/310037

**Authors:** John Rippe, Justin H. Baumann, Daphne N. De Leener, Hannah E. Aichelman, Eric B. Friedlander, Sarah W. Davies, Karl D. Castillo

**Affiliations:** Department of Marine Sciences, University of North Carolina at Chapel Hill, 3202 Murray Hall, Chapel Hill, NC, USA.; Department of Statistics and Operations Research, University of North Carolina at Chapel Hill, 318 Hanes Hall, Chapel Hill, NC, USA.; Curriculum for Environment and Ecology, University of North Carolina at Chapel Hill, 3202 Murray Hall, Chapel Hill, NC, USA.

**Author notes:** Department of Biological Sciences, Old Dominion University, 110 Mills Godwin Life Sciences Building, Norfolk, VA, USA. Department of Biology, Boston University, 5 Cummington Mall, Boston, MA, USA.

**Keywords:** Coral reef, calcification, Caribbean, Florida Keys, sclerochronology, climate change, global warming, ocean acidification

## Abstract

Through the continuous growth of their carbonate skeletons, corals record invaluable information about past environmental conditions and their effect on colony fitness. Here, we characterize century-scale growth records of inner and outer reef corals across ~200 km of the Florida Keys Reef Tract (FKRT) using skeletal cores extracted from two ubiquitous reef-building species, *Siderastrea siderea* and *Pseudodiploria strigosa.* We find that corals across the FKRT have sustained extension and calcification rates over the past century but have experienced a long-term reduction in skeletal density, regardless of reef zone. Notably, *P. strigosa* colonies exhibit temporary reef zone-dependent reductions in extension rate corresponding to two known extreme temperature events in 1969-70 and 1997-98. We propose that the subtropical climate of the FKRT may buffer corals from chronic growth declines associated with climate warming, though the significant reduction in skeletal density may indicate underlying vulnerability to present and future trends in ocean acidification.

## INTRODUCTION

As the most thoroughly monitored coral reef ecosystem in the wider Caribbean, the Florida Keys Reef Tract (FKRT) has unfortunately become a paradigm for the severe decline of coral reefs across the region throughout the last four decades. Following the near extirpation of acroporids in the 1970s and 1980s, a further decline in stony coral cover on the order of 40% since 1996 has driven an ecological shift on the FKRT towards greater dominance of octocorals, macroalgae and sponges (Miller, Bourque, & Bohnsack, 2002; Ruzicka et al., 2013; Somerfield et al., 2008). This deterioration of the stony coral community has been attributed primarily to chronically warming waters (Causey, 2001; Manzello, 2015), acute high and low temperature stress events (Colella, Ruzicka, Kidney, Morrison, & Brinkhuis, 2012; Kemp et al., 2011; Lirman et al., 2011), and widespread disease (Porter et al., 2001; Precht, Gintert, Robbart, Fura, & Van Woesik, 2016).

Yet, sustained reductions in coral cover have been unevenly skewed towards outer reef environments, highlighting a unique cross-shelf distinction on the FKRT (Ruzicka et al., 2013). Inner patch reefs along the seaward boundary of Hawk Channel, a 10 m deep channel running 2-3 km offshore of the Florida Keys archipelago, maintain 15-17% stony coral cover despite their proximity to the highly variable water conditions associated with Florida Bay (Ruzicka et al., 2013). By comparison, outer reef sites, which are bank-barrier reefs located 8-9 km offshore along the edge of the shelf predominantly immersed in the clear waters of the Florida Current, have been reduced to ≤5% coral cover and continue to experience significant mortality of important reef-building coral species (Ruzicka et al., 2013).

Depending on geographic context, two competing theories are often proposed to explain differences in ecology and resilience of corals at inner versus outer reef sites. In some cases, observations of higher coral cover (Lirman & Fong, 2007; Thomson & Frisch, 2010), greater colony size and growth rates (Manzello, Enochs, Kolodziej, & Carlton, 2015; Soto, Muller Karger, Hallock, & Hu, 2011), elevated bleaching resistance (Barshis et al., 2013; Palumbi, Barshis, Traylor-Knowles, & Bay, 2014), and stable growth trajectories (Castillo, Ries, & Weiss, 2011; Castillo, Ries, Weiss, & Lima, 2012) on nearshore reefs suggest that consistent exposure to a highly variable environment preconditions resident coral populations to better cope with ocean warming and the increasing frequency of high temperature stress events. However, conflicting reports of reduced coral cover (De’ath & Fabricius, 2010), degraded thermal tolerance (Carilli, Norris, Black, Walsh, & McField, 2010), and slower coral growth rates (Cooper, De’Ath, Fabricius, & Lough, 2008) suggest that exposure to high levels of suspended sediments and nutrients associated with local human development reduces the fitness of nearshore corals. It is more likely that the evolutionary or acclimatory advantage gained by living in a variable thermal environment co-occurs with the negative impacts of terrestrial runoff, and the balance of these factors determines the relative condition of the reef community.

Sclerochronology, or the use of coral skeletal cores to examine historic trends in growth, provides a useful tool in diagnosing reef health in space and time, and therefore can reveal differences in the sensitivity of sampled reef areas to their changing environments. Early such studies from the upper Florida Keys report long-term growth trends of the *Orbicella* species complex, drawing qualitative comparisons between skeletal extension rates and anthropogenic factors related to human development in south Florida (Hudson, 1981; Hudson, Hanson, Halley, & Kindinger, 1994). More recent analysis of *Orbicella faveolata* from the same region reveals that variations in skeletal density and extension rates over time correlate significantly with the Atlantic Multidecadal Oscillation index (Helmle, Dodge, Swart, Gledhill, & Eakin, 2011). Additionally, baseline calcification rates were found to be significantly greater at an inner reef site of the upper Florida Keys relative to a nearby outer reef site (Manzello et al., 2015). These findings support the premise that there may be a physiological growth advantage for corals living in inner reef environments of the FKRT.

Notably, no sclerochronology studies on the FKRT thus far have found significant long-term trends in extension or calcification rates, which deviates from reports on other reef systems. In the Pacific, Andaman Sea and Red Sea, for example, calcification rates of *Porites* spp. and *Diploastrea heliopora* have declined alongside rising ocean temperatures over the past three decades (Cooper et al., 2008; De’ath, Lough, & Fabricius, 2009; Tanzil, Brown, Tudhope, & Dunne, 2009). Similarly, multiple studies on the Belize Mesoamerican Barrier Reef System (MBRS) have revealed a century-scale decline in the extension rates of *Siderastrea siderea* and *Pseudodiploria strigosa*; although, the trend in this case varied based on proximity to shore and the spatial scale of investigation (Baumann et al., 2018; Castillo et al., 2012). Along a single inner-outer reef transect on the southern MBRS, forereef colonies of *S. siderea* were found to exhibit a long-term decline in extension rates, while those from nearshore and backreef environments maintained stable growth trajectories (Castillo et al., 2011; Castillo et al., 2012). The authors suggest that this trend may arise due to local water quality dynamics or due to lower resilience of forereef corals to rising ocean temperatures. Future investigation, however, conducted at the scale of the entire reef system, revealed a contrasting pattern in which declining extension rates were observed only for colonies of *S. siderea* and *P. strigosa* at nearshore sites, but not for colonies farther from shore (Baumann et al., 2018). Such complexity of coral growth trajectories throughout the Belize MBRS reflects the delicate balance between the historical advantage of residing in a variable nearshore environment and the deteriorating conditions associated with terrestrial runoff and continual ocean warming.

Here, we assess growth trajectories of two abundant and ubiquitous Caribbean reef-building coral species (*Siderastrea siderea* and *Pseudodiploria strigosa*) from four inner-outer reef transects spanning ~200 km of the FKRT. Comparing long-and short-term patterns in extension, density and calcification, we demonstrate that these two coral species have largely sustained extension and calcification rates throughout the FKRT, but have experienced a chronic reduction in skeletal density over the past century. These results suggest that a subtropical climate may buffer corals on the FKRT from warming-induced declines in extension and calcification rates, as is observed in other reef systems, but we propose that declining density may indicate underlying vulnerability to changing carbonate chemistry on the FKRT.

### MATERIALS & METHODS

#### Study design

In May 2015 and 2016, skeletal cores were collected from colonies of the reef-building corals, *Siderastrea siderea* and *Pseudodiploria strigosa*, from four pairs of inner-outer reef sites spanning the Florida Keys Reef Tract (Fig. 1). From south to north, inner reef sites include W Washerwoman (WW), Cheeca Rocks (CR), Basin Hill Shoals (BH) and Bache Shoals (BS). Outer reef sites include E Sambo (ES), Alligator Reef (AR), Carysfort Reef (CF) and Fowey Rocks (FR). In total, 39 *S. siderea* cores and 31 *P. strigosa* cores were collected from 3 to 5 m depth. Cores were extracted using a CS Unitec Model 2 1335 0010 hydraulic drill affixed with hollow extension rods and a 5 cm diameter wet diamond core bit. At each of the eight sites, five healthy colonies of each species were selected randomly for coring. In some cases, less than five colonies of *S. siderea* and *P. strigosa* were collected, either because the dive team was unable to locate five colonies of sufficient size at certain sites or because coring efforts were halted due to inclement weather. Notably, large colonies of *P. strigosa* were relatively rare at the four outer reef sites, whereas colonies of *S. siderea* were ubiquitous across the entire reef tract (Table 1).

**Table 1.**
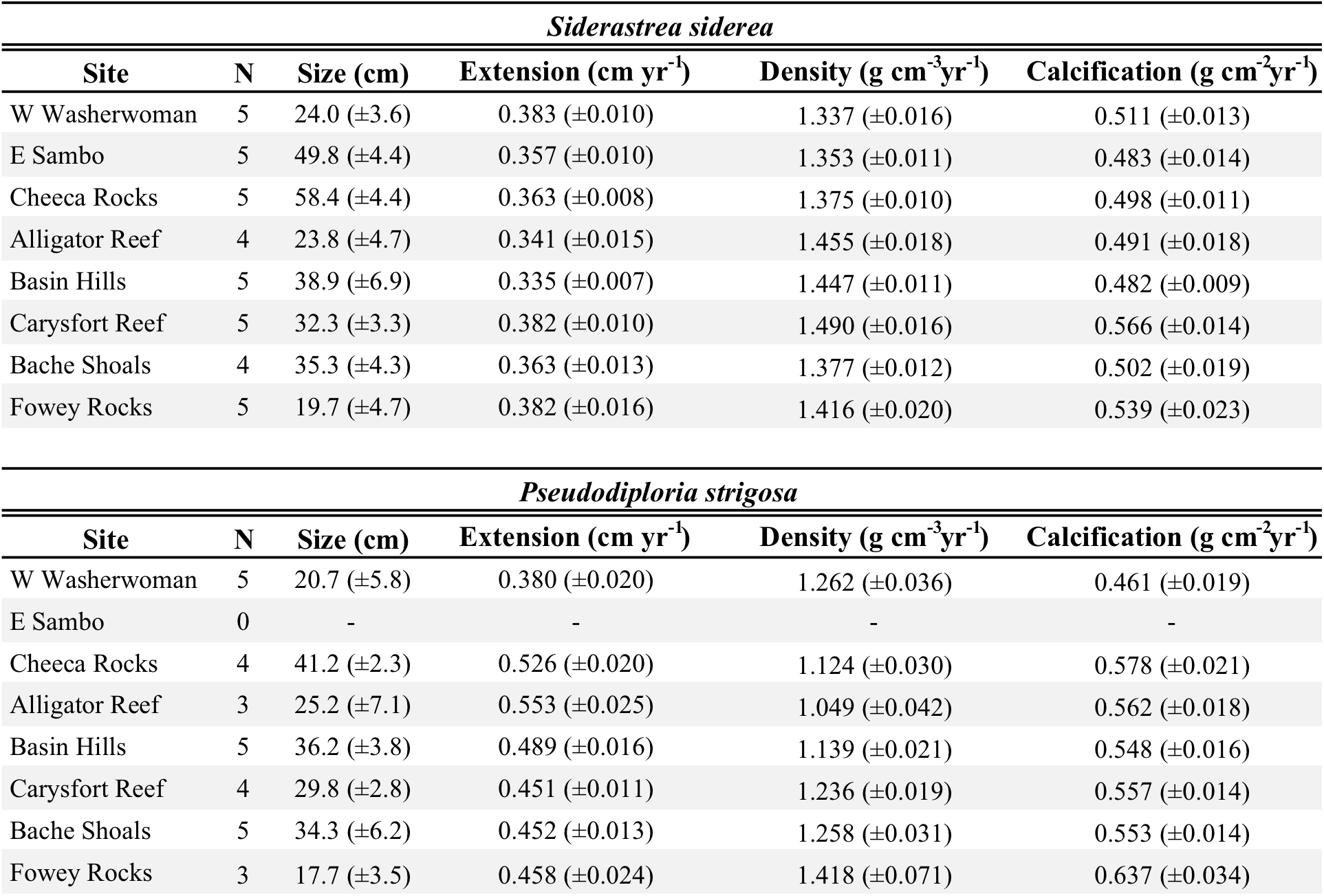
Summary of growth parameters for *Siderastrea siderea* (top) and *Pseudodiploria strigosa* (bottom). Site-wide averages of colony size (estimated by core length), annual extension rate, density and calcification rate

| *Siderastrea siderea* |  |  |  |  |  |
| --- | --- | --- | --- | --- | --- |
| Site | N | Size (cm) | Extension (cm yr<sup>-1</sup>) | Density (g cm<sup>-3</sup>yr<sup>-1</sup>) | Calcification (g cm<sup>-2</sup>yr<sup>-1</sup>) |
| W Washerwoman | 5 | 24.0 (±3.6) | 0.383 (±0.010) | 1.337 (±0.016) | 0.511 (±0.013) |
| E Sambo | 5 | 49.8 (±4.4) | 0.357 (±0.010) | 1.353 (±0.011) | 0.483 (±0.014) |
| Cheeca Rocks | 5 | 58.4 (±4.4) | 0.363 (±0.008) | 1.375 (±0.010) | 0.498 (±0.011) |
| Alligator Reef | 4 | 23.8 (±4.7) | 0.341 (±0.015) | 1.455 (±0.018) | 0.491 (±0.018) |
| Basin Hills | 5 | 38.9 (±6.9) | 0.335 (±0.007) | 1.447 (±0.011) | 0.482 (±0.009) |
| Carysfort Reef | 5 | 32.3 (±3.3) | 0.382 (±0.010) | 1.490 (±0.016) | 0.566 (±0.014) |
| Bache Shoals | 4 | 35.3 (±4.3) | 0.363 (±0.013) | 1.377 (±0.012) | 0.502 (±0.019) |
| Fowey Rocks | 5 | 19.7 (±4.7) | 0.382 (±0.016) | 1.416 (±0.020) | 0.539 (±0.023) |
| *Pseudodiploria strigosa* |  |  |  |  |  |
| Site | N | Size (cm) | Extension (cm yr<sup>-1</sup>) | Density (g cm<sup>-3</sup>yr<sup>-1</sup>) | Calcification (g cm<sup>-2</sup>yr<sup>-1</sup>) |
| W Washerwoman | 5 | 20.7 (±5.8) | 0.380 (±0.020) | 1.262 (±0.036) | 0.461 (±0.019) |
| E Sambo | 0 | - | - | - | - |
| Cheeca Rocks | 4 | 41.2 (±2.3) | 0.526 (±0.020) | 1.124 (±0.030) | 0.578 (±0.021) |
| Alligator Reef | 3 | 25.2 (±7.1) | 0.553 (±0.025) | 1.049 (±0.042) | 0.562 (±0.018) |
| Basin Hills | 5 | 36.2 (±3.8) | 0.489 (±0.016) | 1.139 (±0.021) | 0.548 (±0.016) |
| Carysfort Reef | 4 | 29.8 (±2.8) | 0.451 (±0.011) | 1.236 (±0.019) | 0.557 (±0.014) |
| Bache Shoals | 5 | 34.3 (±6.2) | 0.452 (±0.013) | 1.258 (±0.031) | 0.553 (±0.014) |
| Fowey Rocks | 3 | 17.7 (±3.5) | 0.458 (±0.024) | 1.418 (±0.071) | 0.637 (±0.034) |

**Figure 1.**
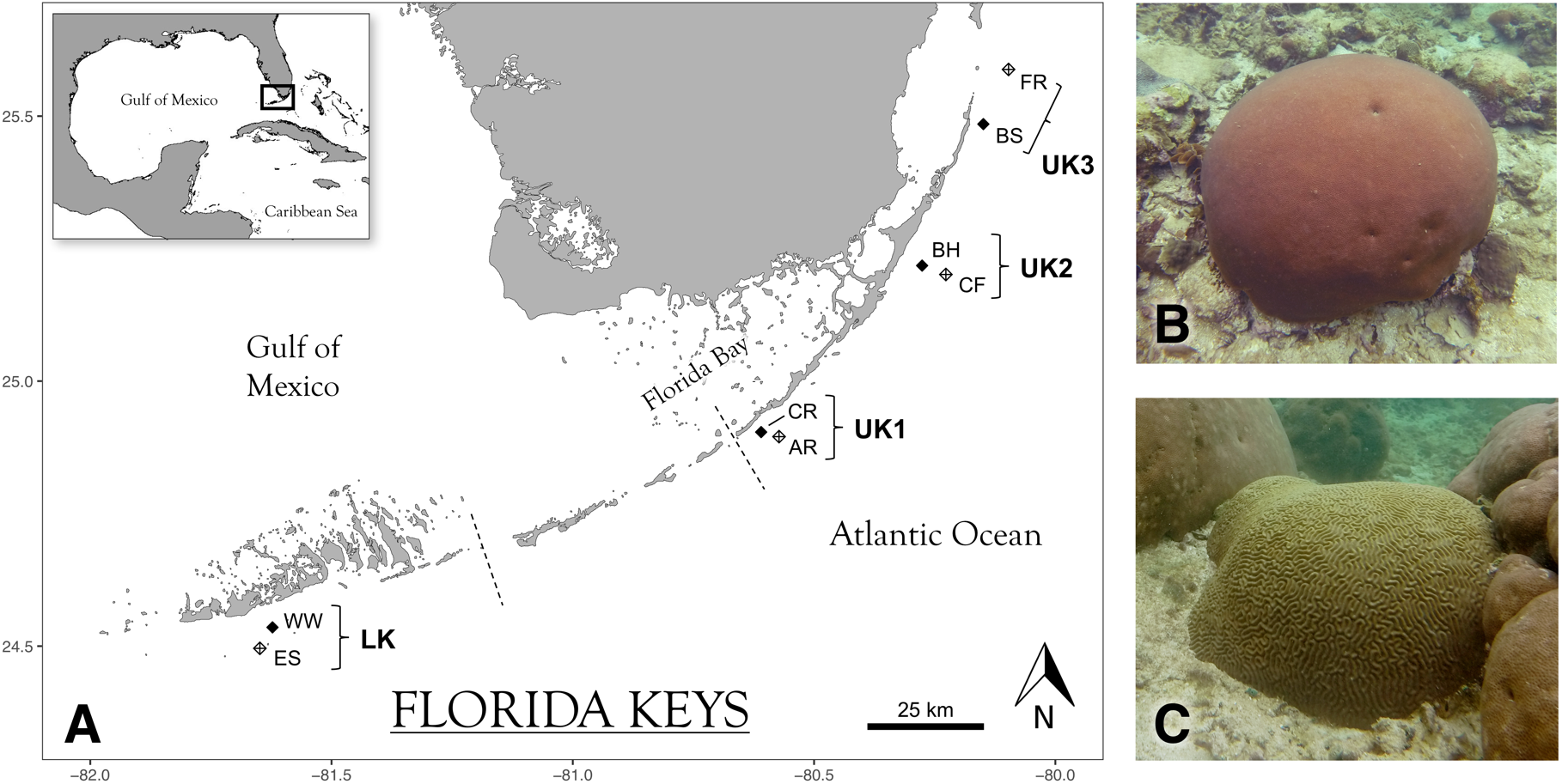
Map of sampling sites (A) and study species (B, C). Solid and crosshatch diamonds represent inner and outer reef sites, respectively. Dashed lines depict the approximate boundaries between the Lower, Middle and Upper Keys. Site and transect abbreviations are as follows: Lower Keys (LK) – West Washerwoman (WW), Eastern Sambo (ES); Upper Keys 1 (UK1) – Cheeca Rocks (CR), Alligator Reef (AR); Upper Keys 2 – Basin Hills (BH), Carysfort Reef (CF); Upper Keys 3 (UK3) – Bache Shoals (BS), Fowey Rocks (FR). The two study species are pictured on the right: *Siderastrea siderea* (top) and *Pseudodiploria strigosa* (bottom).

Cores were extracted from the vertical growth axis of each colony and were at maximum 0.7 m in length, encompassing up to 137 and 89 years of growth for *S. siderea* and *P. strigosa,* respectively. After extraction, a concrete plug was inserted and secured in the drilled hole with Z-Spar^®^ underwater epoxy to protect the colony from erosion and further physical damage. The collected cores were then stored in capped PVC tubes filled with 100% ethanol (EtOH) and transported to the University of North Carolina at Chapel Hill where they were air dried in preparation for sclerochronology development.

#### Sclerochronology development

To assess coral skeletal growth histories, all cores were scanned using X-ray computed tomography (CT) on a Siemens Biograph CT scanner at the Biomedical Research Imaging Center, University of North Carolina at Chapel Hill. Coral cores were oriented lengthwise in rows of 4 to 5 on the scanning table, and equipment parameters were set to 120 kV, 250 mAs and 0.6 mm slice thickness with images reconstructed at 0.1 mm increments using the H70h “Very Sharp Spine” window. All images were exported from the scanner as DICOM files, which were then 3-dimensionally reconstructed using the open-access Horos v2.0.2 medical image viewing software. High- and low-density bands were visualized using a 10-mm thick ‘Mean’ projection oriented as a rectangular prism through the center of each core (Supp. Fig. S4).

All boundaries between semiannual density bands were delineated manually and three sets of linear transects were drawn down the length of the cores using the Region of Interest (ROI) tool in Horos (Supp. Fig. S4). Each set of transects was drawn within the exothecal space between corallite walls in order to standardize density measurements and to avoid aberrant density spikes in areas where the transect may otherwise have crossed a high-density corallite wall. Density and calcification measurements are therefore lower than would be expected if all features of the skeletal architecture were taken into account. Importantly, it has been shown that individual colonies may vary in their timing of high- and low-density band deposition due to intraspecific differences in tissue thickness and morphology (Barnes & Lough, 1993, 1996; J. Carricart-Ganivet, Vásquez-Bedoya, Cabanillas-Terán, & Blanchon, 2013; Taylor, Barnes, & Lough, 1993). Thus, to approximate a consistent time standard between cores, we begin all chronologies at the top of the first fully deposited density band beneath the band of terminal growth. Additionally, because cores were collected in subsequent years, the most recent year of growth (2015) was not included for cores collected in 2016 in order to keep the beginning of chronologies uniform throughout the study.

By-pixel density measurements were extracted from linear transects and average density was calculated for each semiannual high- and low-density band. Following previously established protocol (DeCarlo et al., 2015), nine coral standards of known density were included in every scanning session to convert density measurements from CT Hounsfield units to g cm^−3^. Average density of each standard was assessed in Hounsfield units using Horos and a standard curve was created for all cores scanned in the corresponding session (Supp. Fig. S5). Linear extension was measured in Horos as the width of each annual density band couplet, and calcification (g cm^−2^) was calculated as the product of density and linear extension.

#### Statistical analysis

Mean growth parameters were calculated for each coral species within each site by averaging annual measurements of extension, density and calcification across time. Additionally, as a coarse representation of colony size, the physical length of each core sample was also measured and averaged within sites (Table 1).

Colony-scale variability in annual extension, density and calcification was evaluated using complementary methods, first via coefficients of variation (CV) and then by testing for significance of spatial autocorrelation in the most recent 30 years of growth (1985-2014). We employ a 30-year threshold in order to compare sufficiently long-term chronologies while also retaining a large majority of sampled cores (58 of 67 samples). All cores shorter than 30 years were not included in this analysis. CV provides a relative measure of variation irrespective of the mean, calculated as the ratio of the standard deviation to the mean 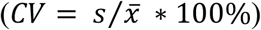.

Interannual CV was quantified for each sampled colony (Fig. 2A). Spatial autocorrelation was evaluated using a permutation-based Mantel test (n=1000), and the resulting correlogram was fitted with a non-parametric correlation spline and 95% confidence interval determined by bootstrapping (n=1000; Fig. 2B; Supp. Fig. S3). The Mantel test was conducted using an increment of 10 km to create 13 uniformly spaced pairwise distance classes. Analyses of spatial autocorrelation were performed in the *ncf* package in R (Bjornstad, 2009; R Core Team, 2017).

**Figure 2.**
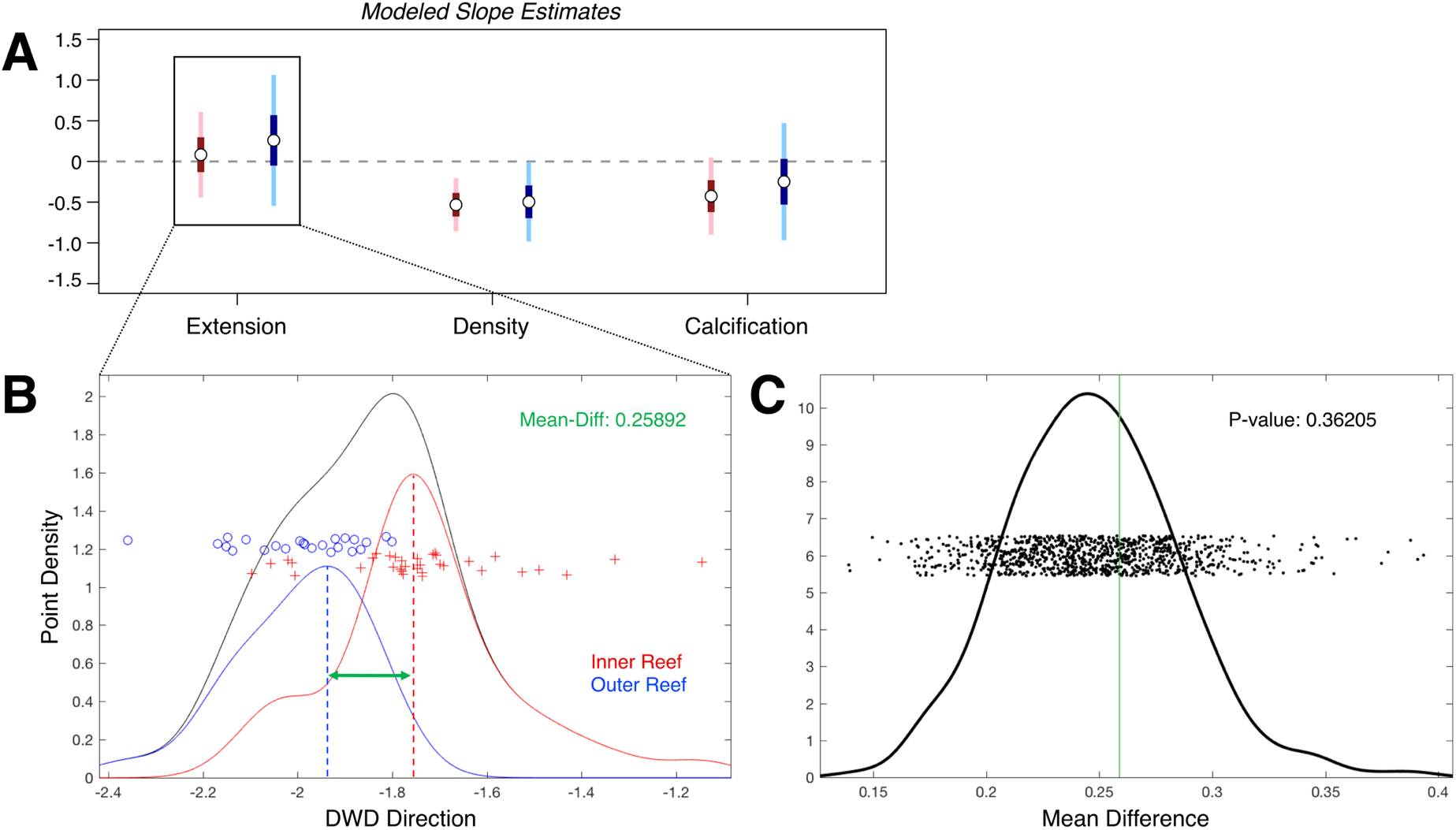
Long-term growth patterns do not differ between reef zones. (A) Modeled estimates of linear trends in extension, density and calcification for the inner (red) and outer reef (blue). Dark- and light-colored bars represent 50% and 95% confidence intervals, respectively. (B) *Extension* chronologies of inner and outer reef cores were also compared using Distance Weighted Discrimination (DWD). Red crosses (inner reef) and blue circles (outer reef) represent the relative position of each core chronology along the DWD axis, and the mean difference between the two groups was calculated. (C) The significance of the difference between the inner and outer reef cores along the DWD axis was evaluated using a DiProPerm test. Each point signifies a mean-difference calculation after each of 1000 permutations of randomly relabeling the data and refitting the DWD direction. The green line denotes the mean difference between the true groups (inner and outer reef) in the data. Curves in (B) and (C) represent the density of points along the x-axes. All analyses pictured were performed on the full dataset including mean-standardized annual measurements of both study species together.

General differences in patterns of coral growth between inner and outer reef zones were assessed preliminarily using the concept of object-oriented data analysis, which treats each coral growth chronology as a data object, or essentially as one data point (see An et al. (2016) for a detailed description). Within this framework, the annual growth measurements of each chronology exist in *d*-dimensional space (specifically 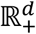), where *d* is the number of years included in the analysis. As in the analysis of spatial autocorrelation, only the most recent 30 years of growth are compared, and all shorter cores are not included for the reasons described above. The vector in this space which best separates the two classes of corals (inner and outer reef) was then computed using a method known as Distance Weighted Discrimination (DWD) (Marron, Todd, & Ahn, 2007). Since the data exists in such high dimension, we expect there to be a direction in M-which separates the two classes almost perfectly regardless of whether a difference actually exists. To remedy this, we conducted a Direction-Projection-Permutation (*DiProPerm*) statistical test (Wei, Lee, Wichers, & Marron, 2016) to evaluate whether the vector separating the two groups appears to be better than one would expect solely due to the high dimensionality of the data. The test is conducted by randomly relabeling the class of each core sample (inner and outer reef) and refitting the DWD direction 1,000 times. A *p*-value is then calculated as the percentage of iterations that are separated better than the DWD direction of the original data.

Lastly, temporal trends in growth were assessed in two steps, first using linear mixed effect (LME) modelling, generally following the statistical protocol of Castillo et al. (2011), and second using generalized additive modelling to capture short-term fluctuations in growth. In order to account for the hierarchical nature of the dataset, for both modelling approaches standardized annual values of linear extension, density and calcification measured within each core were treated as the units of observation, while the cores themselves were treated as sampling units and were incorporated as random effects. Consistent with Castillo et al. (2011), the variable *Year* in all LME models was centered to minimize correlation between random slopes and intercepts, and the residual correlation structure of individual cores was described using an autoregressive moving-average model of order (*p,q*). An interaction term between *Reef Zone* and *Year* is included as a model predictor in order to compare linear trends in growth between reef zones (Appendix 1).

Basic generalized additive models (GAM) for extension, density and calcification were implemented in the *mgcv* package in R (R Core Team, 2017; Wood, Pya, & Säfken, 2016), incorporating an adaptive smoothing spline of *Year* as a fixed effect predictor and *Core* as a random effect. Adaptive smoothing allows the degree of smoothing to vary with the covariates to be smoothed and is therefore well suited to capture rapid fluctuations in growth. The smoothing basis (*k* = 25) was selected following the protocol recommended by Wood (2017), whereby *k* was increased progressively (*k* = 5, 10, 15, 20, 25) until the effective degrees of freedom stabilized at a value sufficiently lower than *k* – 1. Once the models were fitted to the data, time intervals of significant change were computed using the first derivative of the fitted trend spline, following the methods of Bennion, Simpson, and Goldsmith (2015). In short, a finite differences approximation of the first derivative is calculated at fixed time points along the model prediction with an associated 95% confidence interval. Where the confidence interval of the derivative curve excludes 0 (i.e., zero slope), we conclude that significant change in growth is observed at that time point. These intervals of significant change are indicated on all GAM plots as thick green (increasing) and red (decreasing) segments.

## RESULTS

### Coral growth trajectories on the Florida Keys Reef Tract

Coral cores of *Siderastrea siderea* and *Pseudodiploria strigosa* were collected from four pairs of inner and outer reef sites on the Florida Keys Reef Tract: three site pairs spanning the southern, middle and northern Upper Keys region (UK1, UK2 and UK3, respectively) and one site pair in the Lower Keys region (LK; Fig. 1). Long-term linear trends and short-term fluctuations of annual extension, density and calcification rates were evaluated for both coral species using linear mixed effects and generalized additive modelling, respectively. We then used a combined Distance-Weighted Discrimination (Marron et al., 2007) and *DiProPerm* (Wei et al., 2016) statistical methodology to compare multi-dimensional patterns of growth between inner and outer reef colonies.

Long-term linear trends reveal that skeletal density of both species significantly decreases through time, while extension and calcification rates are neither increasing nor decreasing (Fig. 2A). We find no evidence to suggest that temporal patterns in extension (*p* > 0.05, Fig. 2B-C), density (Supp. Fig. S1A) or calcification (Supp. Fig. S1B) differ significantly between inner and outer reef sites at the scale of the FKRT (*p* > 0.05 for all parameters). Generalized additive model results also demonstrate little apparent difference in long- or short-term trends between reef zones in either species and, in fact, highlight the considerable colony-level variation in growth through time (Fig. 3). Even accounting for short-term fluctuations, model predictions explain only 5.58% and 4.70% of the deviance in the chronologies of annual extension rate for *S. siderea* and *P. strigosa*, respectively.

**Figure 3.**
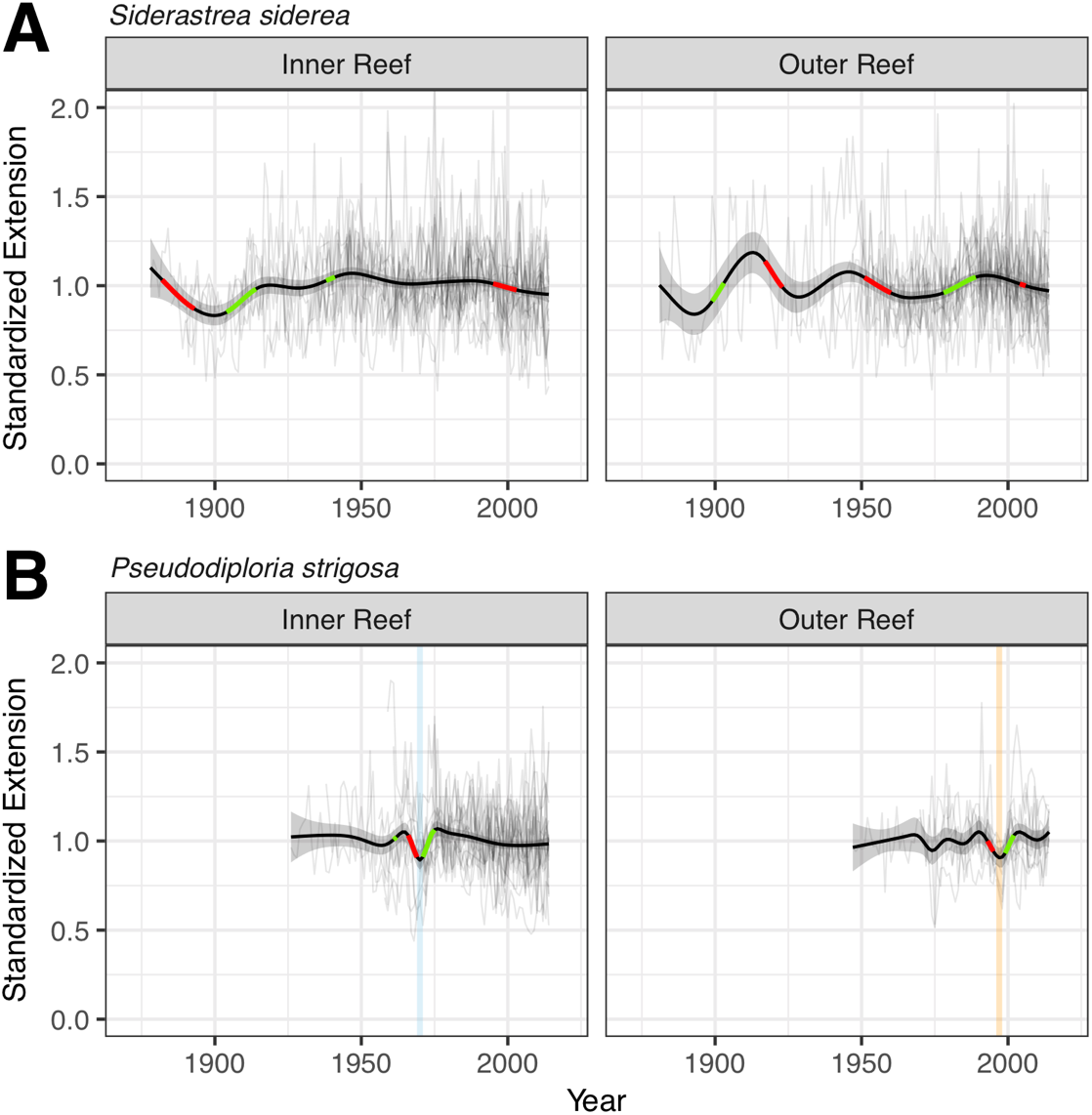
Sensitivity to extreme thermal events differs between species and reef zones. Generalized additive model predictions (±2 SE) overlay individual core chronologies of standardized annual extension (light gray), organized by reef zone and species. Green and red segments denote regions of the curve that are significantly increasing and decreasing, respectively. The light blue and orange vertical lines indicate two previously reported major bleaching events in the FL Keys: 1969-70 cold water (blue) and 1997-98 warm water (orange).

We also compared growth chronologies of each species between the four inner-outer reef site pairs, highlighting the along-shore variability in coral growth trends along the FKRT (Supp. Fig S2). Notably, the extension rate of both species declined significantly in the Lower Keys transect (LK) during the 1997-98 mass bleaching event and had still not fully recovered by 2014. Declining extension in *S. siderea* at the northern Upper Keys transect (UK3) drives a corresponding reduction in calcification since 2001. Additionally, calcification of *P. strigosa* at the middle Upper Keys transect (UK2) has declined significantly since 1980.

### Extreme temperature events differentially impact Pseudodiploria strigosa

Generalized additive model results reveal the short-term response of *P. strigosa* to two documented extreme temperature events on the FKRT. In 1969-70, the extension rates of inner reef *P. strigosa* were depressed in association with a cold-water bleaching event (Hudson, Shinn, Halley, & Lidz, 1976) (Fig. 3B). Similarly, in 1997-98, the extension rates of outer reef *P. strigosa* were depressed in association with a Caribbean-wide warm-water bleaching event (Causey, 2001) (Fig. 3B). *Siderastrea siderea* did not demonstrate the same reduction in extension in association with extreme temperature events at the scale of inner and outer reef zones (Fig. 3A).

### Mean growth and variability within sites

To compare baseline growth parameters between all sampling sites, we calculated site-wide mean rates of extension, density and calcification for each species (Table 1). On average, extension and calcification rates were significantly greater for *Pseudodiploria strigosa*, and skeletal density was greater for *Siderastrea siderea* throughout the FKRT (*p* < 0.001 for each parameter). Mean extension of *S. siderea* was relatively consistent across all sites on the FKRT, while extension of *P. strigosa* was greatest at the two sites within the middle Upper Keys transect (UK2; i.e., Cheeca Rocks and Alligator Reef) and lowest at W Washerwoman (Lower Keys). Between inner and outer reef sites within each transect, skeletal density of both species was generally greater on the outer reef with only one exception – within the southern Upper Keys transect (UK1), density of *P. strigosa* colonies was greater on the inner reef (Cheeca Rocks; 1.124 ± 0.030) than on the outer reef (Alligator Reef; 1.049 ± 0.042). Similar to extension, site-wide averages of annual calcification rates were relatively consistent for *S. siderea*, but varied considerably between sites with no discernible pattern for *P. strigosa.* Mean calcification of *P. strigosa* was greatest at Fowey Rocks (0.637 ± 0.034) and lowest at W Washerwoman (0.461 ± 0.019).

Annual extension and calcification rates were found to vary substantially within and between all sampled colonies, regardless of their geographic proximity. Average interannual coefficients of variation (CV) in extension for each core were 20.5% (10.7 – 33.0%) and 18.0% (9.8 – 33.9%) for *S. siderea* and *P. strigosa,* and in calcification were 20.1% (11.6 – 35.0%) and 18.6% (11.1 – 29.7%) for *S. siderea* and *P. strigosa,* respectively. By comparison, interannual variability in density was considerably lower, with an average CV of 6.8% (3.2 – 17.1%) and 12.1% (5.1 – 20.2%) for *S. siderea* and *P. strigosa*, respectively (Fig. 4A). Additionally, spatial autocorrelation between standardized annual extension, density and calcification of both species was assessed for the most recent 30 years of data (i.e., 1985-2014). Between the three growth parameters, none of the Mantel *r* statistics calculated for each of 13 uniformly spaced distance

**Figure 4.**
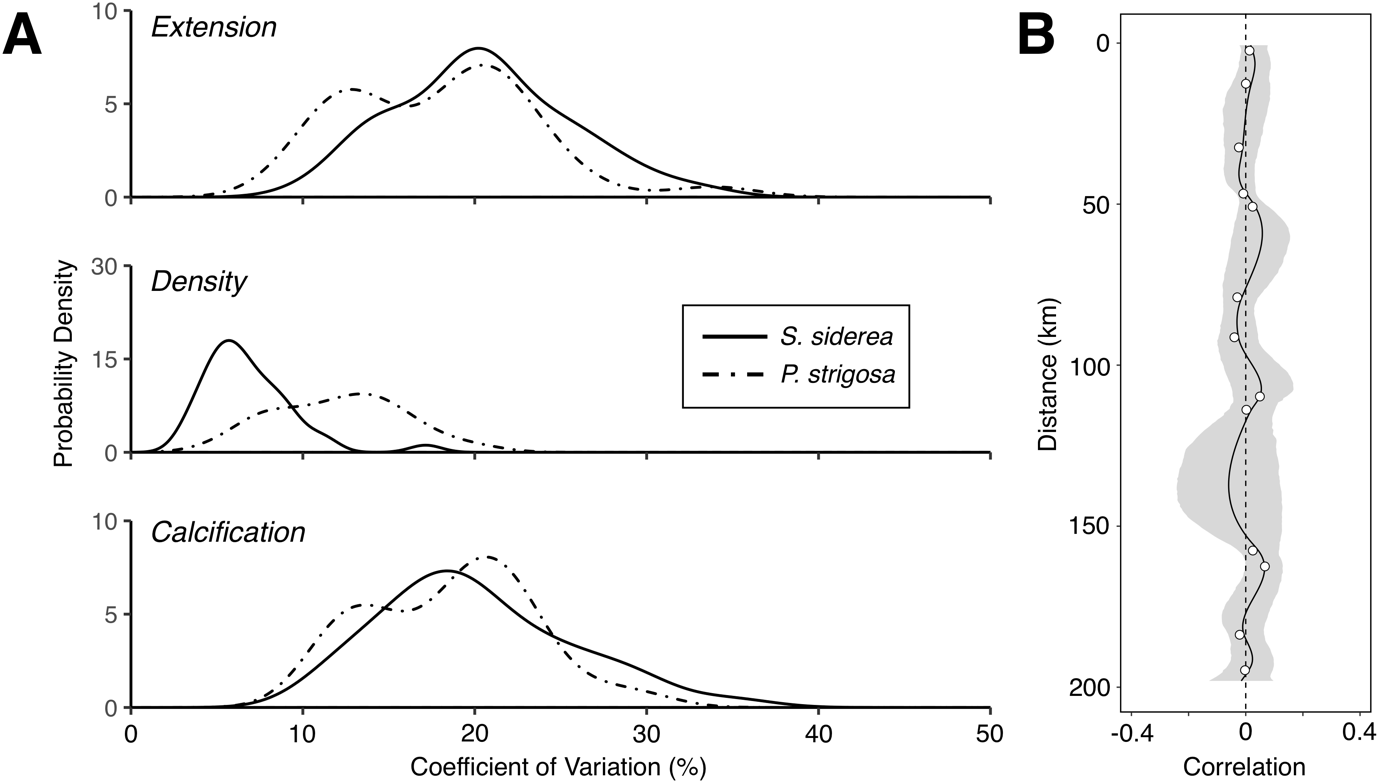
Substantial colony-level variation in annual growth. (A) Probability density plots showing the interannual variation in extension, density and calcification within each core. Coefficient of variation is represented along the x-axis, and the solid and dashed lines depict the density plots for *S. siderea* and *P. strigosa* cores, respectively. (B) Annual extension for the period 1985-2014 was tested for spatial autocorrelation among all cores (*S. siderea* and *P. strigosa).* Open circles represent non-significant Mantel correlation values for 13 uniformly-spaced distance classes within the dataset, determined by permutation test (n = 1000). A nonparametric spatial correlation spline is plotted with a 95% confidence interval determined by bootstrapping (n = 1000). Standardized growth measurements were used to account for differences in mean growth between species and colonies.

classes spanning the FKRT were found to be significant after Bonferroni correction, suggesting no evidence of spatial correlation between nearby colonies (Fig. 4B; Supp. Fig. S3).

## DISCUSSION

This study reveals that the contrasting trends in coral cover and resilience previously observed between inner and outer reefs of the Florida Keys Reef Tract (FKRT) do not translate to clear reef zone differences in long-term growth for *Siderastrea siderea* and *Pseudodiploria strigosa* at the scale of the entire FKRT system. Rather, linear modelling indicates that long-term trends of all growth parameters are virtually indistinguishable between inner and outer reef colonies. Multi-dimensional DWD and *DiProPerm* analyses corroborate this result, offering no evidence that annual growth differs significantly between reef zones in the most recent 30 years (Fig. 2B-C).

The long-term growth trends observed here largely reflect those previously reported for *Orbicella faveolata* from the Upper Keys, with extension and calcification remaining stable and density declining significantly over time (Fig. 2A; (Helmle et al., 2011). However, the lack of reef zone differences contradicts previous observations from the Belize MBRS, which demonstrate variable declines in extension rates based on proximity to shore (Baumann et al., 2018; Castillo et al., 2012). As hypothesized by Helmle et al. (2011), it is possible that the disparity between growth patterns on the FKRT and the MBRS may be attributed to the subtropical nature of the Florida Keys, in that the conditions on the FKRT have not yet crossed a critical temperature optimum that would lead to significant declines in coral growth on a reef zone scale.

Previous research has shown that coral extension and calcification rates are correlated with temperature in a parabolic fashion. At moderate temperatures, coral growth accelerates with increasing temperature (Lough & Barnes, 2000); however, once a thermal optimum is reached, calcification declines with further warming (Castillo, Ries, Bruno, & Westfield, 2014; Jokiel & Coles, 1977). Recent theory supports the existence of such an optimum based on physiological factors known to regulate biomineralization (Wooldridge, 2013). Moreover, a comprehensive *in situ* record of *Porites* growth rates from the Great Barrier Reef reflect this pattern temporally, with an extended period of increasing calcification until 1990 followed by a drastic decline throughout the modern era of climate warming (De’ath & Fabricius, 2010).

A key difference between thermal conditions on tropical and subtropical reef environments is the degree, frequency and duration of extreme temperature events, all of which are factors known to weaken coral health and increase the likelihood of bleaching (Manzello, Berkelmans, & Hendee, 2007). Using the Belize MBRS for comparison, previous work has shown that outer reef sites experienced on average 20.0-40.1 days per year of recorded average temperatures above the local bleaching threshold of >29.7°C (Aronson, Precht, Toscano, & Koltes, 2002) between 2003-2012 (Aronson et al., 2002; Baumann et al., 2016). During the same time period, a permanent monitoring station at Molasses Reef (MLRF1), an outer reef site on the FKRT, recorded only 13.2 days per year with average temperatures >29.7°C (National Data Buoy Center). Moreover, if we consider the metric found to correlate best with bleaching occurrence specifically on the FKRT (i.e., number of days annually >30.5°C; (Manzello et al., 2007), we find that MLRF1 recorded on average only 7.3 days per year above this locally-derived bleaching threshold during the same time period of comparison.

Thus, high-latitude reefs that experience relatively mild summer temperatures may still fall beneath the thermal optimum beyond which coral growth is expected to decline. In accordance with this notion, previous work from western Australia found that *Porites* colonies from two southern sites (i.e., highest latitudes) have exhibited an increase in calcification rates with warming over the last century, while those from more tropical regions have shown no change (Cooper, O’Leary, & Lough, 2012). In addition, a recent study conducted in Bermuda, the northernmost reef system in the Atlantic basin, predicts that calcification rates of two dominant corals may in fact increase with moderate increases in ocean temperature over the next century, though only under conservative warming scenarios (Courtney et al., 2017). Similarly, *S. siderea* and *P. strigosa* colonies sampled here appear able to sustain baseline extension and calcification rates through present day, regardless of their reef zone origin and despite persistent warming on the FKRT (Kuffner, Lidz, Hudson, & Anderson, 2015; Manzello, 2015).

However, exceptions to this pattern occur when extreme temperature events punctuate the prevailing subtropical climate on the FKRT. As has been reported extensively, anomalously warm SSTs are increasing in frequency worldwide, causing the near annual recurrence of major coral bleaching events (Van Hooidonk, Maynard, & Planes, 2013). Moreover, corals on the FKRT are occasionally faced with intrusions of anomalously cold water from the neighboring Florida Bay. Florida Bay is a wide, shallow embayment extending south from the mainland peninsula of Florida and west behind the shelter of the Florida Keys (Fig. 1). Bay waters are subjected to extreme seasonal fluctuations in temperature and salinity, as well as elevated suspended sediment and nutrient concentrations associated with freshwater outflow from the Everglades ecosystem (Boyer, Fourqurean, & Jones, 1999). Regional circulation models show consistent tidal exchange and long-term net transport of water from Florida Bay into Hawk Channel through tidal passes primarily in the Middle Keys (Smith & Lee, 2003). Under average weather conditions, this water exchange has a relatively marginal impact on reefs beyond the boundary of the Middle Keys (Szmant & Forrester, 1996). However, during severe winter cold fronts that draw polar air masses across the region, pulses of bay water can reach surrounding inner reef areas of the Upper Keys and, in some cases, cause catastrophic mortality in the reef community (Colella et al., 2012; Kemp et al., 2011; Lirman et al., 2011).

The coral growth records presented here indicate that two particular extreme temperature events induced short-term reductions in extension rates of *P. strigosa* at the scale of entire reef zones (Fig. 3B). The first instance was a cold-water bleaching event in 1969-70, which reportedly caused 80-90% coral mortality at Hens and Chickens Reef, an inner patch reef in the Upper Keys (Hudson et al., 1976). Long-term growth trends indicate the impact of this event spanned inner reef sites across the FKRT, causing a significant drop in the extension rate of *P. strigosa* colonies at these sites. A similar growth response is observed in association with the severe 1997-98 warm-water bleaching event, during which extension rates of *P. strigosa* colonies at outer reef sites were significantly depressed. This pattern suggests that, although extensive bleaching was reported throughout the FKRT in 1997-98, outer reef areas were particularly impacted by this event, echoing the continued decline of stony coral communities on the outer reef since 1998 (Ruzicka et al., 2013). Some have shown that higher turbidity associated with inner reef areas acts to attenuate UV radiation through the water column, thereby reducing coral susceptibility to bleaching (Morgan, Perry, Johnson, & Smithers, 2017). Additionally, a number of coral genera have demonstrated capacity to mitigate the negative effects of thermal stress via heterotrophy (Grottoli, Rodrigues, & Palardy, 2006). One or both of these factors may have lessened bleaching severity or accelerated the recovery of inner reef corals.

Interestingly, a number of other major bleaching events are not evident in the long-term growth records, namely the 2004-05 warm-water and 2009-10 cold-water events, both of which caused extensive coral bleaching and mortality on the FKRT (Colella et al., 2012; Manzello et al., 2007; Wagner, Kramer, & Van Woesik, 2010). The absence of clear reductions in extension rate at the reef zone scale suggests that the impact of these events may not have been as widespread or as severe as the 1969-70 or 1997-98 events. Alternatively, inherent spatial variation in bleaching susceptibility and impact may hinder reliable correlation of acute stress events with long term growth records. However, as extreme temperature anomalies become more frequent on the FKRT (Manzello, 2015), repeated exposure to these thermal stress events may begin to push conditions beyond their thermal optima and lead to future reductions in coral extension rates.

The long-term decline in skeletal density observed for both species throughout the FKRT highlights the complexity of the coral growth response to the impacts of climate change. Together, skeletal density and extension control the rate of calcification, or the annual amount of skeleton accreted by the coral, which is important in determining whether coral reefs are in a state of net framework construction or erosion (Eyre et al., 2018). Small changes to reef-wide calcification budgets can have direct implications on habitat function and viability; however, growing evidence indicates that density and extension are affected independently by different parameters of environmental change.

Early research revealed variations in density based on hydraulic energy of the reef setting, such that denser skeletons strengthened coral colonies exposed to higher wave activity (Scoffin, Tudhope, Brown, Chansang, & Cheeney, 1992). Reduced skeletal density has also been attributed to elevated nutrients and poor water quality associated with heavy influence from nearby development (J. P. Carricart-Ganivet & Merino, 2001; Dunn, Sammarco, & LaFleur Jr, 2012; Edinger et al., 2000). Accordingly, we find that average density of both species is significantly lower at inner reef sites at all but one of the cross-shore transects across the FKRT (Table 1), implying that these factors may limit baseline coral density. However, we have no reason to believe that hydraulic energy on the FKRT has increased significantly over the past century, and in fact, water quality has improved throughout the FKRT since 1995 (reduced turbidity and organic carbon) (Briceño & Boyer, 2014), which should stimulate an increase in skeletal density through time.

Rather, we hypothesize that the long-term reduction in coral density may reflect changing carbonate chemistry on the FKRT over the past century. Recent analysis of *Porites* growth from the central Pacific reveals a strong sensitivity of skeletal density, but not extension, to aragonite saturation state (Ω_arag_) and ocean acidification (Mollica et al., 2018). Likewise, density has been shown to decrease from high to low Ω_arag_ along a natural pH gradient in Puerto Morelos, Mexico and in Milne Bay, Papua New Guinea (Crook, Cohen, Rebolledo-Vieyra, Hernandez, & Paytan, 2013; Fabricius et al., 2011). Our findings also reflect those of Helmle et al. (2011), which demonstrated significant correlation between declining trends in skeletal density of *O. faveolata* on the FKRT and modelled Ω_arag_. It is important to note that *in situ* monitoring efforts are beginning to unravel the complex biogeochemical mechanisms driving daily and seasonal fluctuations in carbonate chemistry on the FKRT; however, long-term records of Ω_arag_ in nearshore reef environments are severely limited (estimates of Helmle et al. (2011) are derived from open ocean parameters). Thus, while further research is required to corroborate a direct association, the coral density decline revealed here may provide insight into underlying trends in local carbonate chemistry. Furthermore, all but the most conservative emissions scenario simulated by CMIP5 predict Ω_arag_ on the FKRT to continue declining significantly throughout the 21^st^ century (Okazaki et al., 2017; Van Hooidonk et al., 2013). Therefore, if the observed reduction in density continues, it is likely that coral skeletons will become more porous in the future, potentially making them more susceptible to chemical and biological erosion (Enochs et al., 2015; Enochs et al., 2016; Webb et al., 2017).

Considerable alongshore variability in short-term coral growth trends and in mean growth parameters reveals the degree of environmental heterogeneity that exists at smaller spatial scales along the FKRT. Specifically, a significant reduction in extension rates occurred for both species on the LK transect during the 1997-98 bleaching event, and neither species had recovered by the time of collection, suggesting that conditions in the Lower Keys may be inhibiting coral recovery from a major bleaching event. Water quality reports indicate that nutrient-laden waters originating from agricultural runoff and local sewage discharge reach outer reef areas of the Lower Keys due to wide connections with Florida Bay (Lapointe, Barile, & Matzie, 2004; Lirman & Fong, 2007). Prolonged nutrient enrichment has been shown to depress coral calcification and therefore may hinder recovery from thermal stress (Koop et al., 2001; Marubini & Davies, 1996). Similarly, the extension rate of *S. siderea* colonies at UK3 and calcification rate of *P. strigosa* colonies at UK2 have been declining significantly since 2001 and 1985, respectively. These patterns suggest species-specific responses to deteriorating conditions at these locations. Daily temperature range has been shown to vary significantly along the FKRT and has been correlated with reductions in growth rate between alongshore transplants of *Porites astreoides* in the upper Florida Keys (Kenkel, Almanza, & Matz, 2015). Additionally, spatial heterogeneity in dissolved nutrient or suspended sediment concentrations associated with outflow from Florida Bay may drive alongshore variation in growth trends.

Site-wide averages of growth parameters reveal that the baseline calcification rate for *P. strigosa* is greatest at the northernmost outer reef site (i.e., Fowey Rocks); however, the sampled colonies at this site were the smallest in size (Table 1). During the course of sampling, the dive team encountered numerous larger, older colonies of *P. strigosa* (100+ years) throughout the sampling site, but all had experienced recent mortality and were virtually extirpated, presumably during the 2014-2015 bleaching event (*pers. obs.).* We hypothesize that the colonies which were able to survive this event may be especially well adapted and productive in their environment, and are therefore able to maintain comparably high calcification rates.

This finding, however, highlights a critical implication regarding the use of coral cover versus growth rates as indices of overall reef health. Because only living colonies were sampled for this analysis, the growth trends reported here represent only those individuals that have survived the major mortality events occurring over the past several decades and are therefore distinctly resilient to environmental change. Additionally, in comparison to other species, it is important to note that *S. siderea* is particularly robust to thermal stress, allowing the species to persist ubiquitously throughout the Caribbean basin and as far north as Onslow Bay, North Carolina (34.5°N) (Macintyre & Pilkey, 1969). Its resilience to both cold- and warm-water stress has allowed it to become one of the most abundant species on the FKRT (Florida Fish and Wildlife Conservation Commission, 2016) and explains why the bleaching-associated growth rate reductions observed in *P. strigosa* are not reflected for *S. siderea*. Consequently, coral growth trends do not reflect the health of the entire coral reef community. Subtler environmental changes that may have significant consequences for more susceptible individuals or species might not be fully captured in the skeletal growth records of the those sampled for this study.

Likewise, the high variation in annual growth rates and general lack of coherence between individual colonies of the same species, even those from the same site, may further obscure any subtle growth perturbations. For example, there are numerous instances in which one or more colonies exhibit a spike in growth in the same year that others from the same site exhibit significant growth reductions. This variation at the colony level may be due to genotypic differences that result in differential responses to a similar environment. For example, a genetic crossing experiment performed with *Acropora millepora* demonstrated significant family-based differences in physiological response to temperature stress, providing evidence that intraspecific phenotypic variance occurs in natural populations (Meyer et al., 2009). Moreover, the observed variability in annual growth measurements could also be due to local variations in environmental conditions at the scale of individual colonies. Factors such as irradiance, water flow and colony morphology, for example, have been shown to influence microenvironments immediately surrounding individual coral colonies, and could be expected to alter growth patterns at such small scales (Jimenez, Kühl, Larkum, & Ralph, 2008).

Overall, results of this study suggest that two important reef-building species that are ubiquitous across the FKRT have been able to sustain baseline rates of extension and calcification despite recent bleaching events and chronic ocean warming. This is not to say that corals on the FKRT are necessarily resilient to the current state of environmental change, but it suggests that the local climate may buffer corals from chronic growth declines associated with climate warming, such as those observed on other Caribbean reefs and globally. Moreover, the significant long-term reduction in skeletal density highlights the importance of measuring each component of coral growth to fully understand the past and future trajectories of coral reefs in the modern era of climate change. We posit that declining density may point to the susceptibility of corals to changing carbonate chemistry on the FKRT, and suggest that corals may experience further skeletal weakening in the future. Additional investigation of coral growth trends for other, perhaps more susceptible species, coupled with targeted analysis of environmental correlates is encouraged to provide a more comprehensive understanding of the trajectory of the reef community as a whole.

## ACKNOWLEDGEMENTS

Funding for this work was provided by National Science Foundation award OCE-1459522 to KDC. All collections were authorized under Florida Keys National Marine Sanctuaries permit #FKNMS-2015-023, Biscayne National Park permit #BISC-2015-SCI-0007 and John Pennekamp Coral Reef State Park permits #03241525 and #01111626. We thank Clare Fieseler and Colleen Bove for their assistance in the field. We also thank Matt Phillips for his expertise and assistance in CT scanning.

## AUTHOR CONTRIBUTIONS

KDC conceived and designed this study. JPR, JHB, HEA, SWD and KDC collected core samples in the field. Data analysis was performed by JPR with valuable input from KDC, JHB, DND and EBF. JPR prepared the manuscript with all authors contributing to its final form.

## DATA ACCESSIBILITY

Data tables including all annual measurements of extension rate, density and calcification rate included in this analysis are made available via the BCO-DMO Digital Repository, https://www.bco-dmo.org/project/635863.

## SUPPORTING INFORMATION

Additional supporting information can be found in the online version of this article.

**Appendix S1** Linear mixed effect model diagnostics for all growth parameters

**Table S1** Geographic coordinates and maximum depth at sampling sites organized by transect and reef zone

**Fig. S1** DWD and DiProPerm results for density and calcification

**Fig. S2** Generalized additive model results for all growth parameters organized by inner-outer reef transect and species

**Fig. S3** Spatial autocorrelation profiles for density and calcification

**Fig. S4** Step-by-step core analysis procedure in Horos image viewing software

**Fig. S5** Density calibration curve

